# Functional expression of a Mo-dependent formate dehydrogenase in *Escherichia coli* under aerobic conditions

**DOI:** 10.1101/2023.10.27.564357

**Authors:** Marion Schulz, Anne Berger, Ivan Dubois, Valérie Delmas, Mélodie Cadillon, Madeleine Bouzon, Volker Döring

## Abstract

**Background:** Oxygen tolerant complex metal-dependent formate dehydrogenases hold potential for biotechnological applications.

**Principle Findings:** In this work, we report the functional expression of the complex, molybdenum-dependent soluble formate dehydrogenase encoded by the *fdsGBACD* operon from *Cupriavidus necator* (CnFDH) in *Escherichia coli.* Expression of the operon from plasmids or from a copy integrated in the chromosome enabled growth of an energy-auxotrophic selection strain on formate as sole energy source under aerobic conditions. Growth could be accelerated in turbidostat, leading to a drop of the generation time of 1 hour. While no mutation was found in the operon of evolved isolates, genome sequencing revealed non-synonymous point mutations in the gene *focA* coding for a bidirectional formate transporter carried in all isolates sequenced. Reverting the mutations led to a drop in the growth rate demonstrating the *focA* mutation as principle target of continuous culture adaptation.

**Significance:** A member of the oxygen-tolerant subclass of complex FDH showed stable formate oxidation activity when expressed in the heterologous host *E. coli*, a model organism of biotechnology. The integration of the operon in the chromosome offers the possibility of structure/function studies and activity enhancements through *in vivo* mutagenesis, which can also be applied to CO_2_ reduction in appropriate selection hosts.

## Introduction

Formate dehydrogenases (FDH), a diverse family of enzymes, catalyze the reversible conversion of formate to CO_2_, using NAD(P)/NAD(P)H or other compounds as redox co-substrate [1]. Members of this family can be divided into metal-dependent and metal-independent FDHs. The latter are monomeric proteins that do not contain redox-active centers, they are oxygen-insensitive and depend on NADH as redox cofactor [2]. Due to their simple structures and complete O_2_ tolerance, metal-independent enzymes are the ones mostly used in biotechnological applications, notably for the regeneration of NADH, but also in the reductive sense in electrochemical [3] and photoelectrochemical processes [4]. By contrast, the metal-dependent enzymes contain either a molybdenum or a tungsten atom as part of a pyranopterin guanosine dinucleotide (PGD) cofactor, at least one Fe/S-center and have a complex quaternary structure [5]. Although most of these enzymes are oxygen sensitive, membrane bound and require electron donors/acceptors other than NAD(H), a few O_2_-tolerant, NAD(H)-dependent and soluble enzymes have been found among this class, notably from the metabolically versatile bacteria *Cupriavidus necator* (CnFDH) [6, 7] and *Rhodobacter capsulatus* (RcFDH). The cryo-EM structure of this latter enzyme was solved, providing first insights into their mechanism of catalysis [8].

Most FDHs preferentially catalyze the exergonic oxidation of formate to CO_2_. However, under appropriate thermodynamic conditions, they can reduce CO_2_ to formate [9], thus having the potential to become valuable catalysts in the circular carbon economy: the greenhouse gas CO_2_ is converted to value-added formate, that can be used as hydrogen storage material, as fuel in “Direct Formic Acid Fuel Cells” [10], as a versatile C_1_ synthon for chemical synthesis and as sustainable feedstock for the bioindustry [11, 12]. The different FDH enzyme classes vary in this capacity [13], with the O_2_-sensitive metal-dependent dehydrogenases from anaerobic bacteria like *Acetobacter woodii* [14] being the most active (*k*_cat_ > 500 sec^-1^), and the metal-independent dehydrogenases being the least active catalysts (*k*_cat_< 1 sec^-1^). While examples exist in the literature in which the activity of a metal-independent FDH was enhanced up to 3-fold by site directed mutagenesis [15], it can be assumed that the subclass of O_2_-tolerant, NAD- and metal-dependent enzymes have a higher potential to become the enzyme workhorses of CO_2_ reduction under aerobic conditions. The cytoplasmic FDH purified from *Cupriavidus necator* was shown to catalyze this reaction with a *k_cat_*= 11 sec^-1^ under anaerobic conditions [6]. However, enzyme purification was conducted under fully aerobic conditions demonstrating oxygen tolerance despite the presence of a molybdenum-containing CO_2_-formate redox active site and four [4Fe-4S] centers and one [2Fe-2S] center in the α-subunit (FdsA, 105 kDa), a FMN cofactor for NAD/NADH electron transfer and a [4Fe-4S] center in the β-subunit (FdsB, 55 kDa) and a [2Fe-2S] in the γ-subunit (FdsG, 19 kDa).

While *ex vivo* structural and activity studies with complex FDHs have been conducted in recent years, studies of their activity in a cellular context are scarce. Recently, a Mo-dependent enzyme homologously expressed in *Pseudomonas putida* was shown to be active in a selective context [16]. In this report, we describe the cloning and heterologous plasmid-borne expression of the *fdsGBACD* operon coding for CnFDH in an *E. coli* MG1655 derived energy auxotrophic selection strain. We obtained expression-dependent aerobic growth on formate as sole source of energy and isolated faster growing strain descendants upon evolution in continuous culture harboring genetic background mutations. Sustained formate dependent growth was also obtained when the operon was inserted into the chromosome of an evolved isolate cured from the plasmid. Expression from one copy upon genomic integration stabilizes the construct and will enable long-term strain adaptation and evolution in chosen genetic backgrounds to ameliorate enzyme activity and tolerance to O_2_.

## Results

### Rescue of an energy auxotrophic E. coli strain through formate oxidation by FDH from C. necator

The soluble NAD- and Mo-dependent native formate dehydrogenase from *C. necator* is coded by the *fdsGBACD* operon, with the genes *fdsGBA* specifying the three enzyme subunits and the genes *fdsCD* specifying two chaperones shown to be essential for enzyme activity [17, 18]. FdsD was recently shown to be part of the FdsGBAD heterotetrametric functional unit of the closely related FDH from *Rhodobacter capsulatus* [8]. The operon was amplified by PCR from chromosomal *C. necator* DNA and cloned into plasmid pTRC99a. We chose this vector for its strong inducible tac promoter assuring high operon expression. The resulting plasmid pTRC-CnFDH (pGEN1340) was introduced into an *E. coli* MG1655 strain deleted for the gene *lpd* coding for lipoamide dehydrogenase (strain G5416) yielding strain G5663 (for strain and plasmid description, refer to Table 1). This enzyme, a component of the pyruvate dehydrogenase and the 2-oxoglutarate dehydrogenase complexes, catalyzes electron transfer from pyruvate and 2-oxoglutarate to NAD^+^, respectively. *E. coli* strains lacking lipoamide dehydrogenase activity require, when fed with acetate as sole carbon source, an energy source in addition for growth. In this context, NAD-dependent formate oxidation to CO_2_ can provide the necessary energy, as was shown for the monomeric NAD-dependent Fdh of *Pseudomonas* sp.101 [19]. When the expression of the plasmid-borne *C. necator fdsGBACD operon* was induced by IPTG addition in the culture of strain G5663 [Δ*lpd* pTRC-CnFDH], growth was obtained in mineral medium in the presence of formate (60 mM), acetate (20 mM) and pyruvate (20 mM) (Fig. 1). In contrast, no growth was observed when formate was omitted in the culture medium or in the case the energy auxotroph did not harbor the plasmid pTRC-CnFDH (Fig. 1), demonstrating that the *C. necator* NAD-dependent cytoplasmic formate dehydrogenase is functional when expressed in the *E. coli* host. As previously reported [20], we observed low growth yield on formate (60 mM) and acetate (20 mM) as sole carbon source, which was enhanced through pyruvate addition. Pyruvate when replacing acetate supported sustainable growth on formate as energy source. We speculated that the production of acetate through the action of pyruvate oxidase, catalyzing the oxidative decarboxylation of pyruvate to acetate [21, 22] was responsible for this supporting effect. However, deletion of the gene *poxB* specifying the enzyme did only slightly affect growth, pyruvate still being a growth-enhancing factor (not shown). Acetate might be produced from pyruvate by an activity other than pyruvate oxidase, pyruvate formate lyase (Pfl) being a candidate, even so this enzyme is described inactive in the presence of oxygen. In addition, pyruvate might function as supplementary carbon source through gluconeogenesis and as precursor of several amino acids, while it cannot function as electron donor for growth.

**Fig 1.**
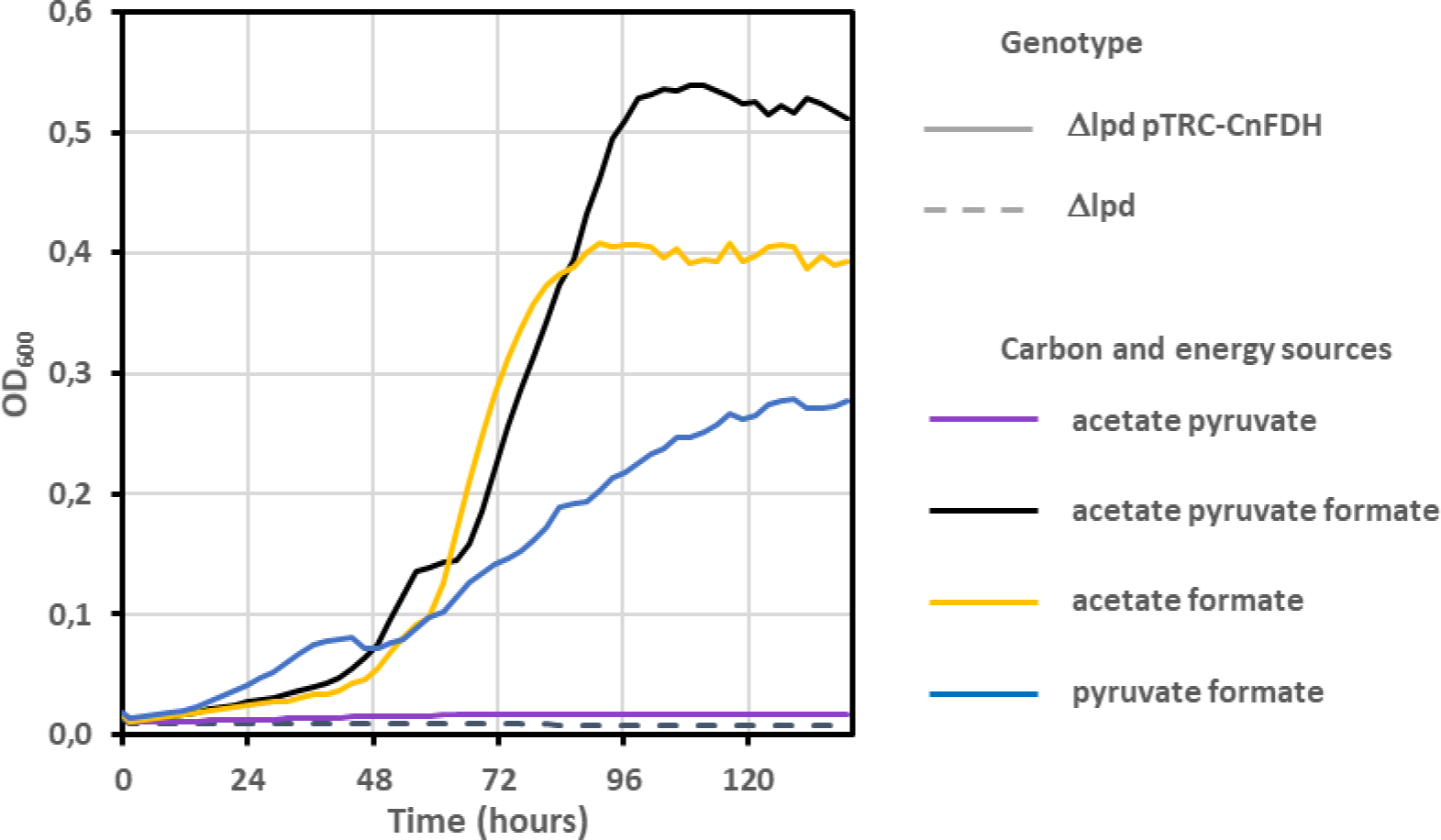
Expression of *C. necator* NAD-dependent formate dehydrogenase allows *E. coli* NADH auxotroph strain Δ*lpd* to use formate as energy source. Strains G5416 (Δ*lpd*) (broken line) and G5663 (Δ*lpd* pTRC-CnFDH) (plain line) were grown at 30°C on mineral MS medium supplemented with the indicated compounds. Concentrations of formate, acetate and pyruvate were 60, 20 and 20 mM, respectively. Growth was recorded with a Bioscreen C plate reader in triplicates.

**Table 1.** Strains and plasmids used in this study.

| Strain | Genotype | Origin/modification |
| --- | --- | --- |
| MG1655 | F<sup>-</sup>, LAM<sup>-</sup>, *rph-1* | CGSC Collection, Yale |
| JW0889 | $\Delta ycaP ::kanR$ | Keio Collection, |
| G5416 | $\Delta lpd$ | Laboratory collection |
| G5663 | $\Delta lpd$<br>pGEN1340 | Transformation of G5416 |
| G5823 | $\Delta lpd focA F7C infC E95K$<br>pGEN1340 | Evolvant of G5663 selected in GM3 under turbidostat regime – UOF1 lineage |
| G5824 | $\Delta lpd focA F7C infC E95K$<br>pGEN1340 | Evolvant of G5663 selected in GM3 under turbidostat regime – UOF1 lineage |
| G5825 | $\Delta lpd focA F7C infC E95K$<br>pGEN1340 | Evolvant of G5663 selected in GM3 under turbidostat regime – UOF1 lineage |
| G5848 | $\Delta lpd focA V97I pps S2F crr G2V fabR L53W$<br>pGEN1340 | Evolvant of G5663 selected in GM3 under turbidostat regime – UOF2 lineage |
| G5849 | $\Delta lpd focA V97I pps S2F crr G2V fabR L53W$<br>pGEN1340 | Evolvant of G5663 selected in GM3 under turbidostat regime – UOF2 lineage |
| G5850 | $\Delta lpd focA F7C cra Y28C$<br>pGEN1340 | Evolvant of G5663 selected in GM3 under turbidostat regime – UOF2 lineage |
| G5876 | $\Delta lpd focA F7C infC E95K$ | Cured G5823 |
| G6234 | $\Delta lpd infC E95K \Delta ycaP ::kanR$<br>pGEN1340 | Transduction G5823 x P1 (Keio JW0889) |
| G6235 | $\Delta lpd pps S2F crr G2V fabR L53W \Delta ycaP ::kanR$<br>pGEN1340 | Transduction G5848 x P1 (Keio JW0889) |
| G6114 | $\Delta lpd$<br>pGEN1393 | Transformation of G5416 |
| G6217 | $\Delta lpd$<br>pGEN1395 | Transformation of G5416 |
| G6272 | $\Delta lpd.focA$ F7C *infC* E95K<br>pGEN1395 | Transformation of G5876 |
| G6280 | $\Delta lpd$ *focA* F7C *infC* E95K<br>pGEN1393 | Transformation of G5876 |
| G6408 | IS10::*fdsGBACD C.n.::kan* | Insertion at IS10 by recombination plasmid pGEN1378 in MG1655 |
| G6435 | $\Delta lpd$ *focA* F7C *infC* E95K IS10::*fdsGBACD C.n.::kan* | Transduction G5876 x P1(G6408) |
| G6504 | $\Delta lpd$ *focA* F7C *infC* E95K<br>pGEN1340 | Transformation of G5876 |
| Plasmid | Description | Origin |
| pTrc99a | IPTG-inducible expression vector, pBR322 origin, *bla<sup>r</sup> lacI<sup>q</sup>* | CGSC Collection, Yale |
| pFDH | constitutive expression pZE21 vector, pBR322 origin, streptomycin<sup>R</sup>, *fdh Pseudomonas sp101* | Addgene (#131706) |
| pFD152 | dCas9 (aTc-inducible), spectinomycin<sup>R</sup>, gRNA cloning site | Depardieu & Bikard (2019) |
| pKI_IS10 | R6K origin, conjugative, chloramphenicol<sup>R</sup> kanamycin<sup>R</sup>, | Wenk *et al.* (2018) |
| pGEN1340 | pTrc99a::*fdsGBACD Cupriavidus necator* native operon | This study |
| pGEN1378 | pKI_IS10::*fdsGBACD Cupriavidus necator* native operon | This study |
| pGEN1393 | pZE21::*fdsGBACD Cupriavidus necator* native operon | This study |
| pGEN1395 | pZE21::*fdhThiobacillus sp. E. coli* adapted synthetic gene | This study |

### Acceleration of formate dependent growth in continuous culture

To accelerate formate-dependent growth, a cell population of NADH-requiring strain G5663 growing in selective medium (formate/acetate/pyruvate, see materials and methods) in the presence of IPTG was subjected to a turbidostat in GM3 continuous culture automatons (see materials and methods). Two independent cultures (UOF1 and UOF2) were launched in parallel. Both cultures were characterized by a short adaptation phase during which the initial generation time dropped rapidly from around 4h40 to stabilize at 3h40. Growth of both cell populations continuously accelerated (Fig. 2) until reaching a plateau with a generation time of 2h15 for both cultures, representing a diminution of about 1h25 as counted from the first stabilized plateau. Three isolates were obtained from each culture and formate dependence of growth verified. Isolate G5823 from UOF1 culture was cured from plasmid pTRC-CnFDH upon serial culture in selective medium supplemented with glucose. Plasmid loss was verified by sensitivity to ampicilline and the absence of PCR amplification of the *fdsGBACD* operon. The cured cells (strain G5876) lost their capacity to use formate as an energy source. Introducing plasmid pTRC-CnFDH in the cured cells restored growth on selective medium, showing that the dependency on FDH-catalyzed formate oxidation for energy supply was maintained during strain evolution (Fig. 3).

**Fig 2.**
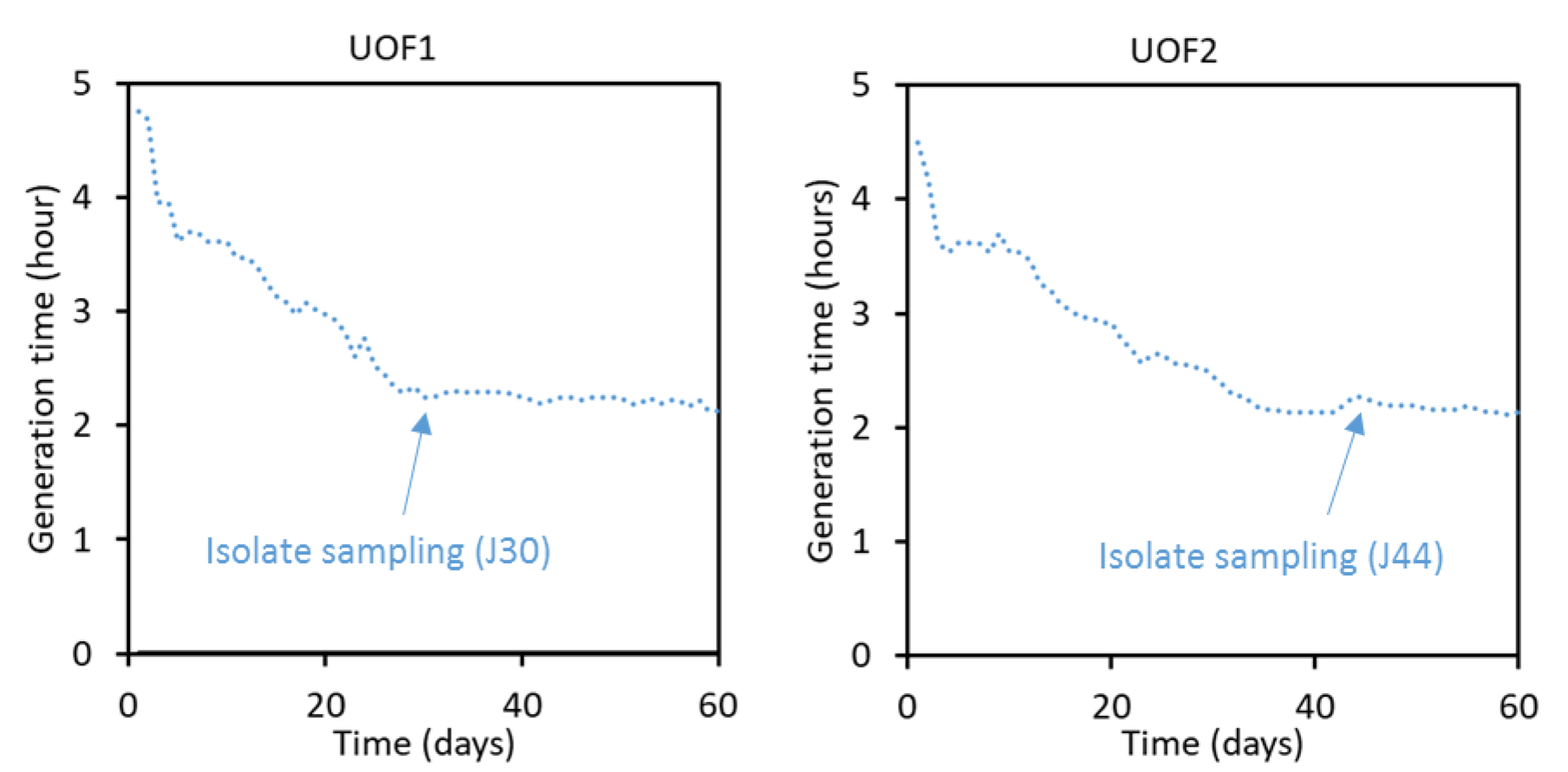
Growth acceleration of NADH auxotroph G5663 bacteria (Δ*lpd* pTRC-CnFDH) in turbidostat. Cells were grown in two independent cultures (UOF1 and UOF2) at 30°C in mineral MS medium supplemented with acetate (20 mM), pyruvate (20 mM) and formate (60 mM) in a GM3 device for 60 days. The time points of isolate samplings are indicated, corresponding to 230 generations in turbidostat for UOF1 and 380 generations in turbidostat for UOF2.

**Fig 3.**
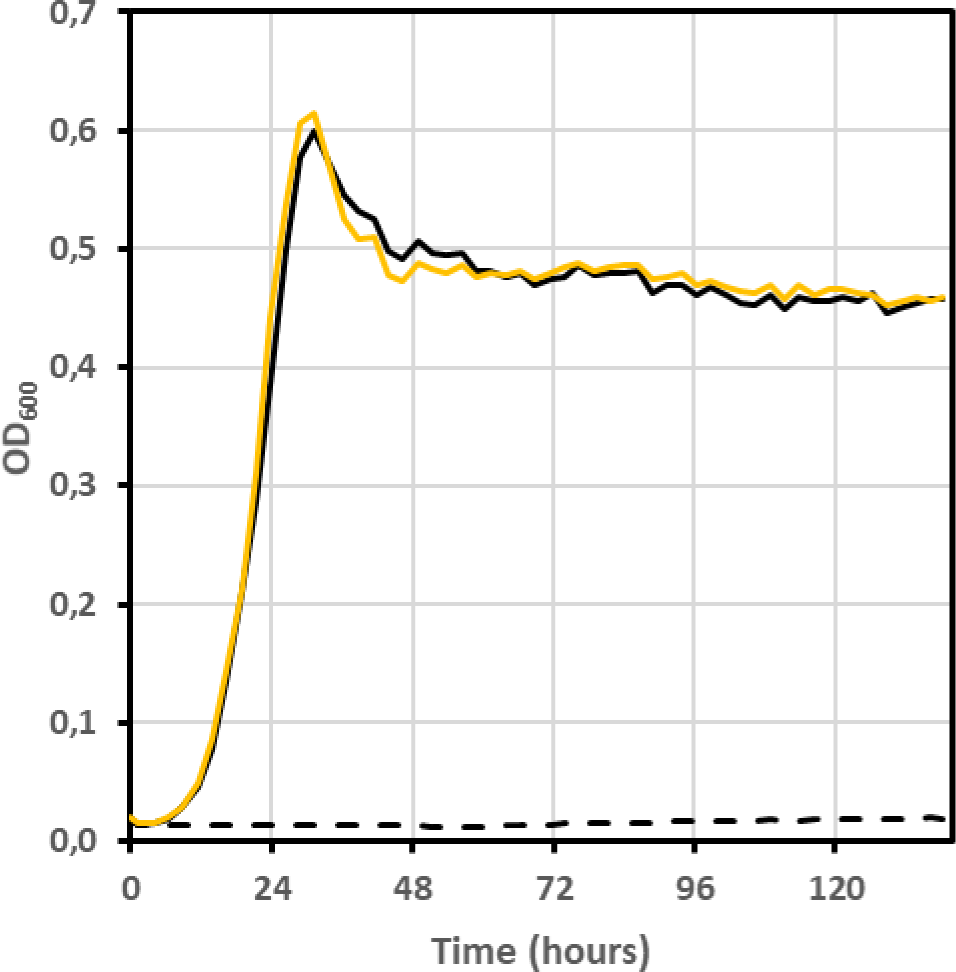
Dependence of evolved isolate G5823 (turbidostat culture UOF1) on the presence of plasmid-borne *C. necator* FDH for growth on formate as energy source. Strains G5823 (black line), G5876 (derivative of G5823 cured from the plasmid pTRC-CnFDH) (black broken line), and strain G6504 (derivative of G5876 transformed with plasmid pTRC-CnFDH) (yellow line) were grown at 30°C on mineral MS medium supplemented with formate (60 mM), acetate (20 mM) and pyruvate (20 mM). Growth was recorded with a Bioscreen C plate reader in triplicates.

To identify adaptive mutations entailing improved growth under selective conditions, we proceeded to whole genome Illumina sequencing of all six isolates and mapped the sequence reads onto the genome (chromosome and plasmid pTRC-CnFDH) of the ancestor strain (see materials and methods). The plasmid pTRC-CnFDH from all six isolates remained unmutated. Sequencing of the genomes of the isolates identified a total of nine point mutations, with eight genes harboring a non-synonymous mutation in their coding region and one mutation affecting an intergenic region (Supplementary Table 1). The only gene found to be affected in all isolates, albeit not carrying the same mutation, was *focA* coding for a bidirectional formate transporter, differing in the changed codon between isolates. Interestingly, one mutation (*focA* F7C) was fixed in isolates obtained from both evolved populations from UOF1 and UOF2 independent cultures. The pH-dependent channel FocA plays an important role in the regulation of intracellular formate concentration, notably during mixed-acid fermentation [23, 24]. We tested the impact of *focA* mutations F7C (isolate G5823) and V97I (isolate G5848) on formate-dependent growth by exchanging the mutated with the wild type *focA* allele in the two isolates. A strong augmentation of the doubling time (3,2x for F7C, 4,3x for V97I) was observed for both derived strains harboring wt *focA* when grown in the formate/acetate/pyruvate test medium (Table 2). FocA regulates formate concentration in the cytoplasm by favoring formate influx or efflux depending on medium pH and the growth phase of cultures in anaerobic environments. In the UOF turbidostats, the population is constantly growing in the logarithmic phase (OD=0,4) at a pH favoring FocA activity in the formate efflux sense (culture medium pH=7.2). Possibly, the mutant variants which arose during evolution impact the fine-tuned regulation of the channel favoring influx or impeding efflux of formate as response to the pressure imposed by the selection.

**Table 2.**
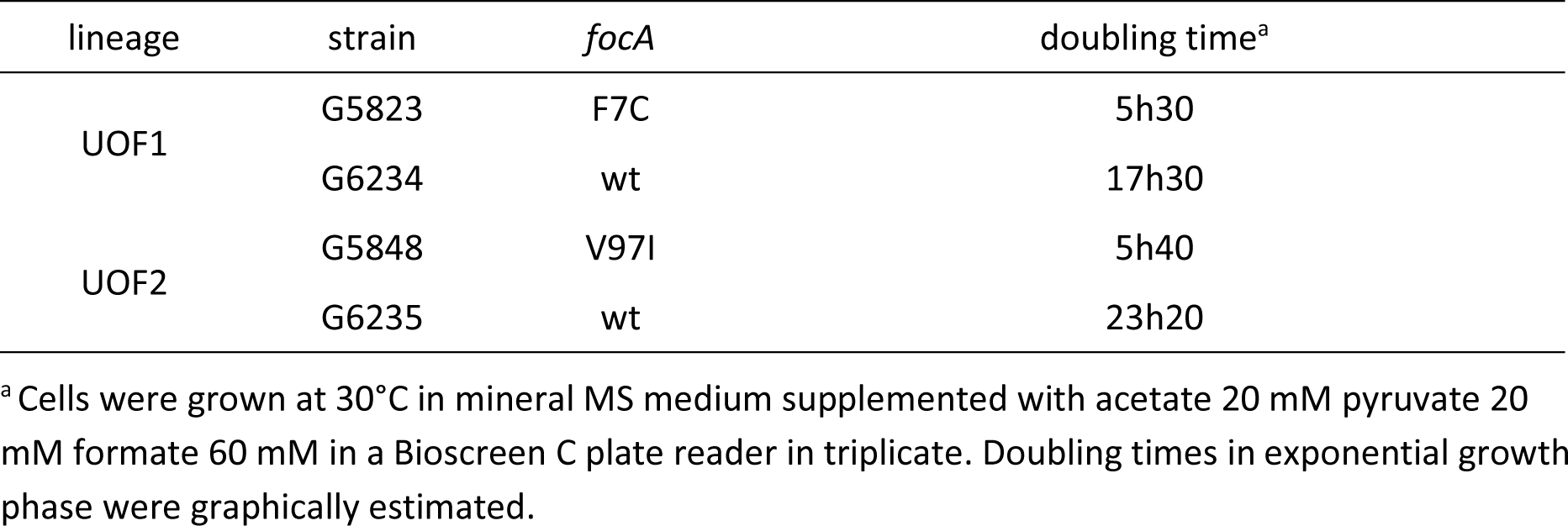
Doubling times of evolved UOF isolates versus derivatives reverted to *focA* wild type.

To test whether the growth rate enhancement effect of the UOF-background was somehow related to the activity of the *C. necator* FDH, we transformed the cured strain G5876 with a pZE21 plasmid (Expressys) for constitutive expression containing the gene for formate dehydrogenase of *Thiobacillus* sp. KNK65MA [25] and compared its growth on formate as sole energy source with the unevolved strain G6272, also harboring the pZE-Ts*fdh* plasmid. In contrast to the *C. necator* enzyme, the enzyme of *Thiobacillus* is monomeric not involving metal centers. Fig. 4 shows that the evolved background also had an enhancing effect on growth under selective conditions when this enzyme was expressed, indicating an adaptation on carbon flux during evolution rather than a regulation of FDH activity related to the structure.

**Fig 4.**
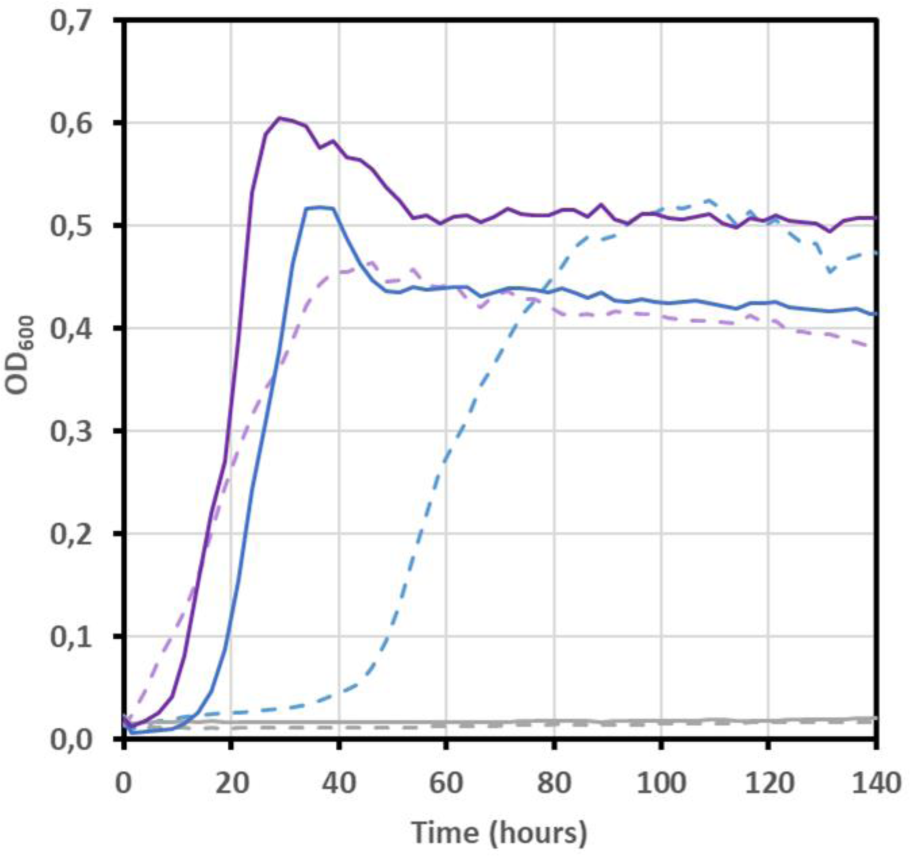
Influence of the genetic background on FDH-dependent growth. Formate-complemented growth of Δ*lpd* strains expressing *C. necator* or *Thiobacillus sp.* formate dehydrogenase was compared in unevolved genetic background of strain G5416 (broken line) and evolved genetic background of strain G5876 (cured derivative of UOF1 isolate G5823) (plain line). Growth of strains G6114 and G6217, derivatives of strain G5416 harboring plasmid pZE::CnFDH (blue broken line) or pZE::TsFDH (purple broken line), respectively, was compared with strains G6280 and G6272, derivatives of strain G5876 likewise harboring plasmids pZE::CnFDH (blue line) or pZE::TsFDH (purple line), respectively. Lack of growth of plasmid-free strains G5416 and G5876 (grey lines) demonstrate the dependence on heterologous FDH activity for cell proliferation under selective conditions. Bacteria were grown on mineral MS medium supplemented with formate (60 mM) acetate (20 mM) and pyruvate (20 mM) at 30°C in a Bioscreen C plate reader in triplicates.

### Chromosomal integration of the complex FDH

To create a platform of stable *C. necator* FDH expression in *E. coli* enabling *in vivo* structure/function studies and the evolution of activity in continuous culture, we integrated the *fdsGBACD* operon in the IS10 site of *E. coli* strain G5876 behind a strong promoter and an RBS following a described protocol [26]. We used the evolved and cured energy auxotrophic strain G5876 originating from culture UOF1 for chromosomal integration to favor our chances to obtain growth on formate. Formate dependent growth was observed for the resulting strain G6435 and the essential implication of the integrated FDH operon demonstrated by CRISPR interference [27]. This method is based on the concomitant expression of a catalytically inactive dCas9 protein and a guide RNA targeting the gene or the operon to be silenced. The inactive dCas9 protein binds – guided by the gRNA - to the promoter or a gene locus near the N-terminus thus interfering with initiation or elongation of DNA transcription by the RNA polymerase. Fig. 5 shows the results for the *C. necator fdsGBACD* operon silenced with three different gRNAs specific for the operon (see materials and methods). Overnight culture samples were serially diluted and dotted on permissive or test plates containing or not the dCas9 expression inducer anhydrotetracycline. Slight growth inhibition was noticed on permissive medium in the presence of the inducer, suggesting residual DNA cleavage activity of dCas9 independent of the presence of a gRNA. On test medium without inducer of dCas9 expression, growth of control cells was observed. Expression of a specific gRNA strongly impeded cell growth under these conditions, demonstrating loss of growth on formate through silencing of the formate dehydrogenase. In the presence of the inducer, virtually no growth was observed, reflecting the deleterious effect of residual dCas9 activity on the Δ*lpd* strains due to their attenuated growth on the test medium.

**Fig 5.**
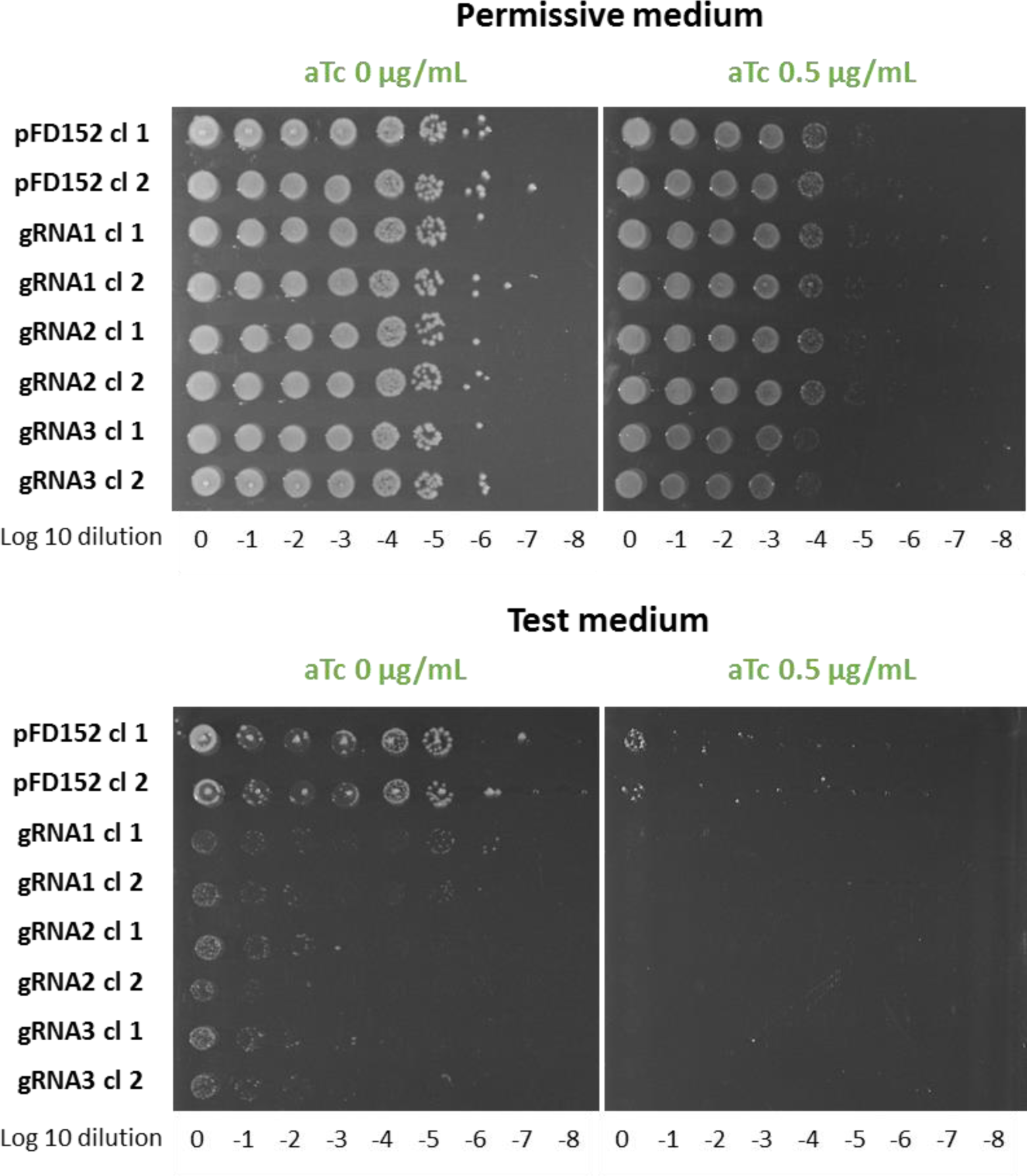
dCas9 silencing of *C. necator fdsGBACD* operon inserted in the chromosome of *E. coli* Δ*lpd* strain. Cells of strain G6435 (Δ*lpd* IS10::*fdsGBACD C.n.)* were grown overnight in permissive medium (MS glucose 0.2% acetate 20 mM), then serially diluted and dotted on semi-solid permissive or test medium (MS formate 60mM acetate 20mM pyruvate 20mM) containing or not the dCas9 inducer anhydrotetracycline (aTc) as indicated. Plates were incubated at 30°C for a maximum of seven days Results are shown for G6435 cells harboring the empty plasmid pFD152 as control, and G6435 cells expressing one of three different gRNAs cloned in pFD152.

Strain G6435 (Table 1) grew with a generation time of 3h30 in the formate/acetate/pyruvate test medium. As expected, induced FDH expression from the multicopy-plasmid pTrc99a supported faster growth on formate (Tgen=1h30) than expression from the single chromosomal copy integrated in strain G6435 (Fig. 6). The linear correlation between formate concentration and growth was demonstrated for the UOF1 isolate G5823 (Fig. 7A) and from its derived strain G6435 (Fig. 7B) for a formate range of 5 to 100 mM.

**Fig 6.**
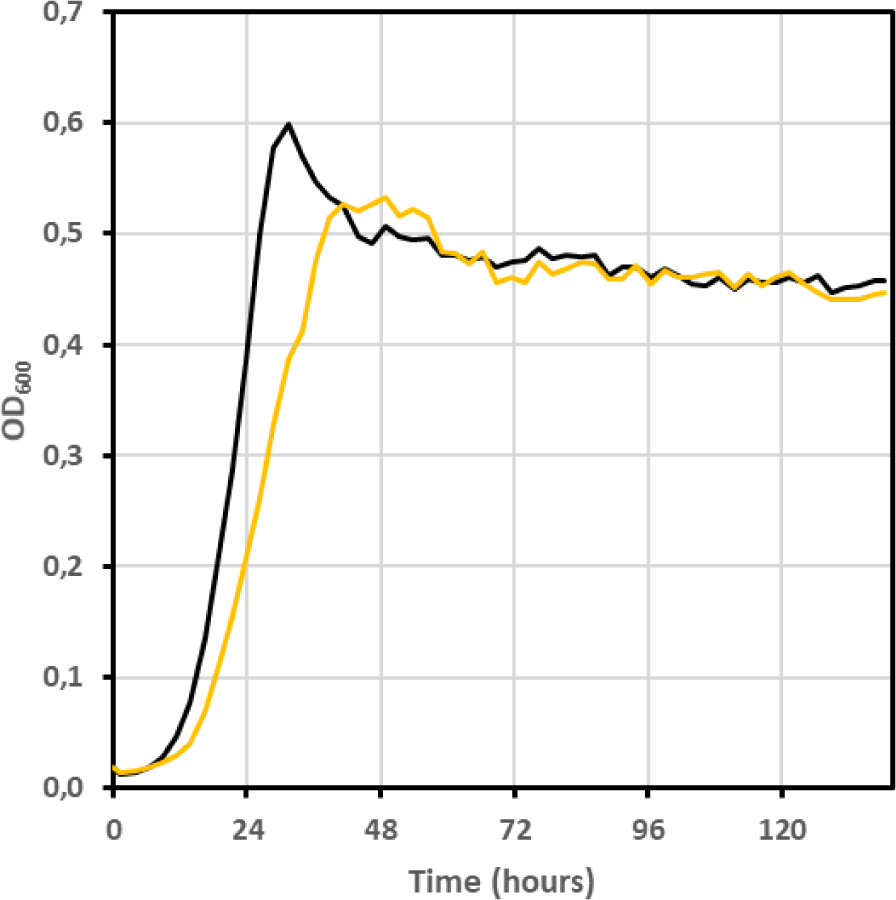
Impact of FDH expression context on growth. Growth of the UOF1 isolate G5823, which harbors plasmid pTRC-CnFDH (black line) was compared with growth of G5823 descendant strain G6435 containing of *C. necator* FDH operon *fdsGBACD* on the chromosome (yellow line). Bacteria were grown on mineral MS medium supplemented with formate (60 mM), acetate (20 mM) and pyruvate (20 mM) at 30°C in a Bioscreen C plate reader in triplicates.

**Fig 7.**
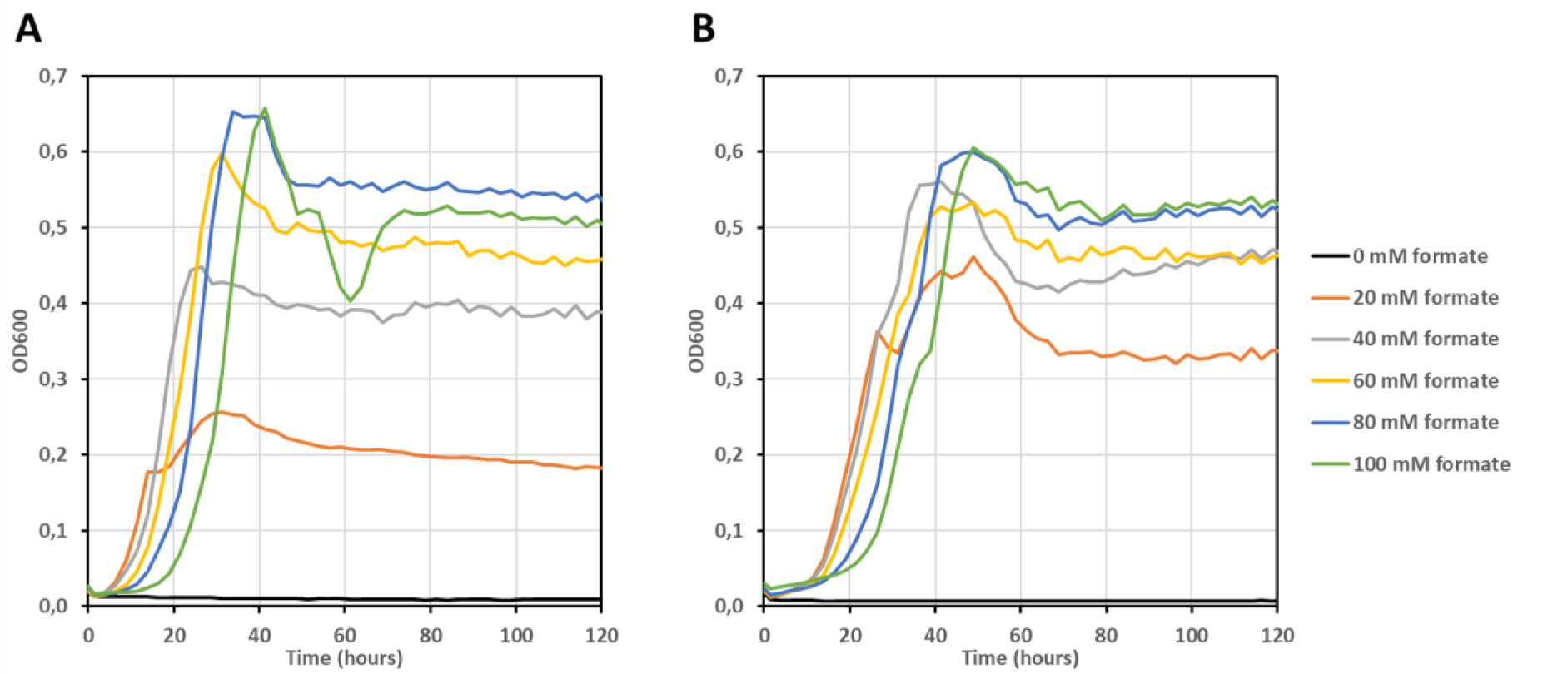
Growth of *C. necator* FDH expressing Δ*lpd* strains depends on formate concentration in the medium. Bacteria from UOF1 isolate G5823 (**A**) and from its derivative strain G6435 (**B**) harboring *C. necator fdsGBACD* operon on the chromosome were grown in mineral MS medium supplemented with acetate 20 mM, pyruvate 20 mM and formate at concentrations ranging from 0 to 100 mM. Overall growth rate was higher for G5823 cells expressing CnFDH from plasmid. The final OD_600_ of both cultures increased with increased formate concentration, the maximum OD being reached at 80 mM for both strains. Experiments were conducted with Bioscreen C plate reader as described in the Material and method section.

We conducted quantitative PCR to compare the expression levels of the enzyme in the two expression formats (normalized for *fdsA*, Table 3). Without surprise, expression of the three genes in the non-evolved plasmid-bearing G5663 context was higher in permissive than in selective medium. For all contexts tested, expression diminished for genes more distant from the promoter, reflecting lower processivity of the RNA polymerase towards the 3’ end of the polycistronic reading frame [28]. The expression level between strain G5663 and its evolved descendant G5823 were comparable for the genes *fdsG* and *fdsA*, but differed for *fdsD*. Given that the mutational analysis of pGEN1340 from strain G5823 did not reveal mutations, the difference observed for *fdsD* expression is not easily explainable. Finally, when comparing the two expression formats, plasmid and chromosome, a lower expression was seen for the G6435 strain which is consistent with the lower growth rate observed when this strain was compared with strain G5823 (Table 3).

**Table 3.**
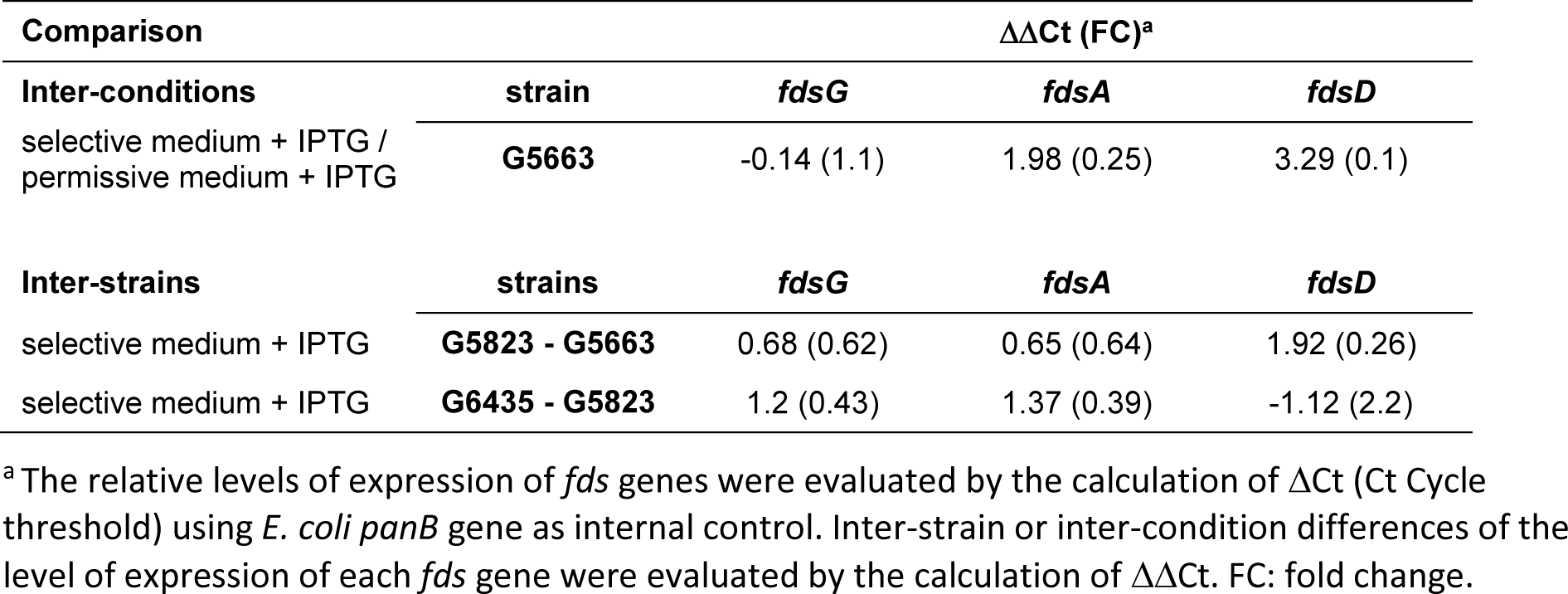
Comparison of the transcriptional levels of the genes *fdsG, fdsA* and *fdsD* by quantitative PCR.

Strain G6435 was subjected to turbidostat evolution in a GM3 and a decrease in generation time of about 1h within 55 days (around 500 generations) was observed for two parallel cultures which reached Tgen=2h32 and 2h14 for cultures OCF5 and OCF6 respectively (Fig. 8).

**Fig 8.**
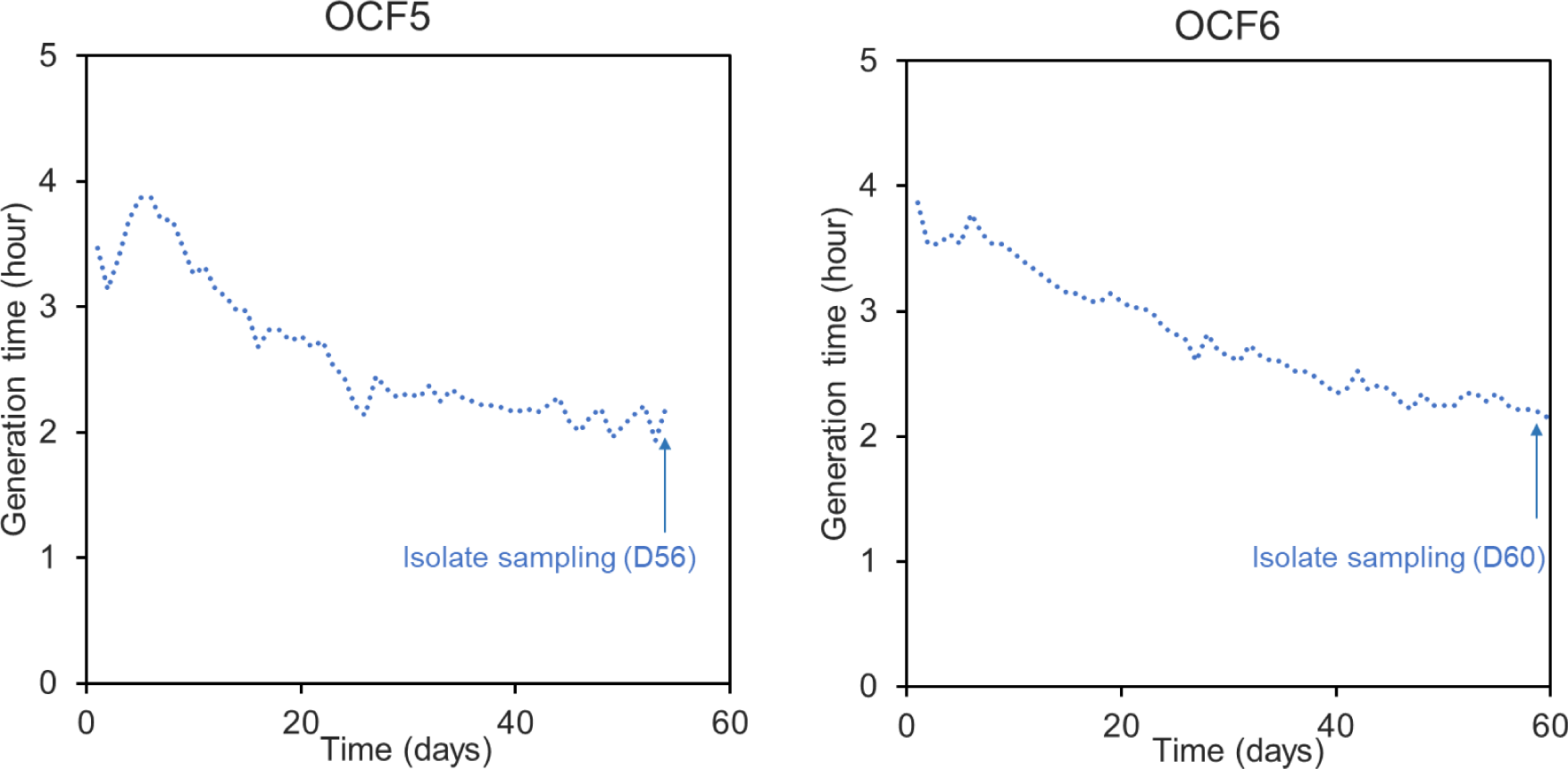
Growth acceleration of NADH auxotroph G6435 bacteria (Δ*lpd* IS10::*fdsGBACD C.n.*) in turbidostat. Cells were grown in two independent cultures (OCF5 and OCF6) at 30°C in mineral MS medium supplemented with acetate (20 mM), pyruvate (20 mM) and formate (60 mM) in a GM3 device for 60 days. The time points of isolate samplings are indicated, corresponding to about 500 generations in turbidostat for both cell populations.

## Conclusion

Complex metal- and NAD-dependent formate dehydrogenases have been identified and studied in recent years. Purification and *in vitro* activity tests under oxic conditions demonstrated oxygen resistance of these enzymes, which was enhanced by stabilizing factors like nitrogen azide [6].

In this study we addressed the question whether such enzymes could stably function *in vivo* under aerobic conditions. An energy auxotrophic *E. coli* strain was used as test system to validate FDH activity for formate oxidation to generate NADH necessary for cell growth [20]. The soluble formate dehydrogenase from *C. necator* expressed from plasmids or from the chromosome supported stable formate-dependent growth in the presence of O_2_. The plasmid-bearing and the chromosomal insertion strains were evolved in continuous culture for faster growth for up to 400 generations, without loss of the selective formate/FDH dependency of the populations. Genomic sequencing revealed adaptive mutations in the genetic background of evolved isolates, while the sequence of the heterologous FDH operon was found unchanged.

The FDH chromosomal insertion construct is of special interest as it provides a stable expression platform not only for continuous culture evolution, but also for *in vivo* site directed or targeted random mutagenesis. In recent years, methods were developed to enable mutagenesis and selection in the same cellular background. Key residues directly involved in the catalytic activity could be identified giving insights into structure/function relationships of these complex enzymes by *in vivo* activity screens avoiding protein overexpression and purification.

As a further perspective, FDH activities could be enhanced for the reductive reaction, using recently constructed formate dependent *E. coli* strains as selection chassis [29, 30]. Efficient enzymatic CO_2_ reduction to formate can be envisioned as an entry point for CO_2_ assimilation for biomass production engineered in initially heterotrophic model strains like *E. coli*. The metal- and NAD-dependent formate dehydrogenases, characterized by their oxygen tolerance and a CO_2_ reduction activity up to 20x higher as compared to non-metal FDHs, are promising candidates for the implementation of synthetic autotrophic growth modes.

## Materials and Methods

### FDH plasmid constructions

The *fdsGBACD* operon (gene IDs 10917038-10917042) coding for the soluble Mo-dependent formate dehydrogenase of *Cupriavidus necator* DSM 13513 (CnFDH) was PCR amplified from genomic DNA using oligonucleotide primers 6125 GAGGTTAATTAAATGCCAGAAATTGCCCCCCAC (fwd) and 6126 ACAGCCAAGCTTTTACTCCAGCATCGCCCGATG (rev). The PCR fragment was gel purified and inserted into plasmids pTRC99a and pKI_IS10 (gift of S. Wenk) using HiFi DNA Assembly Cloning Kit (NEB. pZE21) for Gibson cloning. To obtain integrative plasmid pKI_IS10_CnFDH cloning was conducted in *E. coli* DH5α λpir cells. A version optimized for *E. coli* codon usage of the gene coding for formate dehydrogenase from *Thiobacillus sp*. (AB106890) was synthetized by Twist Biosciences, California and cloned into plasmid pZE21 (Expressys, kan^r^, colE1 origin, tet promoter) using the CPEC protocol [31].

### Strain constructions

The strains used or constructed in this study were all derivatives of the wild type *E. coli* K12 strain MG1655. Their relevant genotypes and filiations are listed in Table 1. The desired genetic contexts were obtained by phage P1-mediated transductions of gene knockouts substituted by antibiotic resistance cassettes according to the method of [32]. Genes of interest were mobilized in the desired recipient cells by co-transduction with closely linked kanamycin markers originating from the Keio *E. coli* knockout collection [33]. Resistance cassettes were removed by flippase reaction after transformation with the plasmid pCP20 . The *fdsGBACD* operon flanked in 3’ by a strong promotor and an RBS was inserted in the chromosomal IS10 site [26] of MG1655 by recombination with plasmid pKI_IS10_CnFDH. The plasmid was transformed into the *E. coli pir*^+^ donor strain ST18 [34] and transferred into recipient strain MG1655 by conjugation. Plasmid integration and subsequent removal of the plasmid backbone were selected as described [26]. The *fdsGBACD* operon was PCR amplified and correct integration verified by sequencing.

### Continuous culture

Evolution experiments in continuous culture were carried out using GM3 fluidic self-cleaning cultivation devices. This device automatically dilutes growing cell suspensions with nutrient medium by keeping the culture volume constant. A continuous gas flow of controlled composition through the culture vessel ensures constant aeration and counteracts cell sedimentation. Twin culture vessels connected with silicon tubing enable the periodical transfer of the evolving culture between vessels and their cleaning upon rinsing with a 5N NaOH solution to remove biofilms [35].

To evolve G5663 cells to faster growth on formate as energy source, a turbidostat regime was programmed. This cultivation regime enables the selection of optimized growth in permissive conditions. Every 10 min, the optical density of the culture is automatically measured and compared to a fixed threshold (OD_600_ value of 0.4). When the measured OD_600_ exceeds the threshold, a pulse of fresh nutrient medium is injected into the culture and the same volume of used culture discarded. The dilutions ensure that the biomass in the vessel remains constant and that the bacteria grow at their maximal growth rate. A preculture of G5663 cells was grown in MS formate (60 mM) acetate (20 mM) pyruvate (20 mM) medium supplemented with IPTG (100 µM) at 30°C to an OD_600nm_ of 0,8 and used to inoculate two independent culture vessels (UOF1 and UOF2) with the same medium composition. Samples of the growing cultures were taken once a week and kept at −80°C. Growth was stopped after 230 (UOF1)and 380 (UOF2) generations and culture samples plated on semisolid MS formate acetate pyruvate medium to obtain isolates from colonies for further analysis. Cultures OCF5 and OCF6 were inoculated in a GM3 device with a preculture of strain G6435 grown in MS formate (60 mM) acetate (20 mM) pyruvate (20 mM) medium at 30°C to an OD_600nm_ of 0,8 and used to inoculate two independent culture vessels. Samples of the growing cultures were taken once a week and kept at −80°C. Growth was stopped after 500 generations and culture samples plated on semisolid MS formate acetate pyruvate medium to obtain isolates from colonies for further analysis.

### Bacterial growth assays

A Microbiology Reader Bioscreen C apparatus (Thermo Fisher Scientific) was used for growth curve recordings. It consists of a thermostatic incubator and a culture growth monitoring device (OD reader). Overnight bacterial cultures were washed once in MS medium and diluted 100-fold in the respective growth medium; 200 µl aliquots of the cell suspensions were distributed into honeycomb 100-wells plates. Each experiment was performed in triplicate. The plates were incubated at 30° or 37°C under continuous agitation. Bacterial growth was followed by recording optical densities at 600 nm every 15 minutes during the indicated time.

### Whole genome sequencing and mutation analysis

Pair-end libraries (2×150 bp) were prepared from 1 µg of genomic DNA of the evolved isolates and sequenced using an MiSeq sequencer (Illumina).

High-throughput sequencing data were analyzed using the PALOMA bioinformatic pipeline implemented in the MicroScope platform [36] (https://mage.genoscope.cns.fr/microscope/home/). In a first step, reads were mapped onto the *E. coli* MG1655 reference (NC_000913.3) using the SSAHA2 package (v.2.5.1). Only unique matches having an alignment score equal to at least half of their length were retained as seeds for full Smith-Waterman realignment [37] with a region extended on both sides by five nucleotides of the reference genome. All computed alignments then were screened for discrepancies between read and reference sequences and a score based on coverage, allele frequency, quality of bases, and strand bias was computed for each detected event to assess its relevance. The mutations (single nucleotide variations and short insertions or deletions) with a score superior to 0.8 with at least five supporting reads were retained.

### Gene silencing

Gene silencing was performed using CRISPRi method [27]. Specific gRNAs were cloned into plasmid pFD152 (gift from Solange Miele) harboring the inducible gene coding for dCas9 and a gRNA cloning sites. Three gRNAs (gRNA1: GGCGCCACGTGGTACAGGTC, gRNA2: AGGTGCATGGCGTGATCACC, gRNA3: AAGCGCTGGCCGAGCATGCG) were cloned into plasmid pFD152 using Golden Gate technique (Bsa I) and tested for silencing of the *fdsGBACD* operon. Plasmids expressing a specific gRNA were transformed into the strain G6435 and serial dilutions of overnight cultures were dotted on large Petri dishes in permissive and test conditions with and without induction of dCas9 by 0.5 µg/ml anhydrotetracycline and incubated for 2 to 7 days at 30°C.

### Expression analysis by reverse transcriptase quantitative PCR (RT-qPCR)

The mRNA levels of *fdsA, fdsG* and *fdsD* genes were determined by RT-qPCR with *panB* as internal standard for expression normalization. Cultures were perfomed in MS mineral medium supplemented with either glucose 0.1 %, acetate 20 mM, pyruvate 20 mM, formate 60 mM, IPTG 0.1 mM and carbenicillin 100 mg/L if necessary or acetate 20 mM, pyruvate 20 mM, formate 60 mM and IPTG 0.1 mM if necessary. Cells were harvested in exponential phase (OD_600_ 0.5-0.6). Total RNA was extracted using the RNeasy Mini Kit (Qiagen, Hilden, Germany) following the manufacturer’s instructions. In summary, 2 volumes of RNAprotect Bacteria Reagent (Qiagen, Hilden, Germany) are added to one volume of bacterial culture. After pelleting the cells, RNA extraction was performed and RNA treated with DNase I (NEB). The quality of the extraction was verified by agarose gel electrophoresis. cDNA were generated through reverse transcription using High capacity cDNA Reverse transcription kit (Applied Biosystems) and their concentrations were determined using Qubit™ ssDNA Assay Kit (Invitrogen). Quantitative real-time PCR experiments were performed with three biological replicates, each analyzed in three technical replicates using the KAPA SYBR FAST kit (Roche). Primer pairs used for amplification of genes *fdsA, fdsG* and *fdsD* and *panB* are listed in the table below:

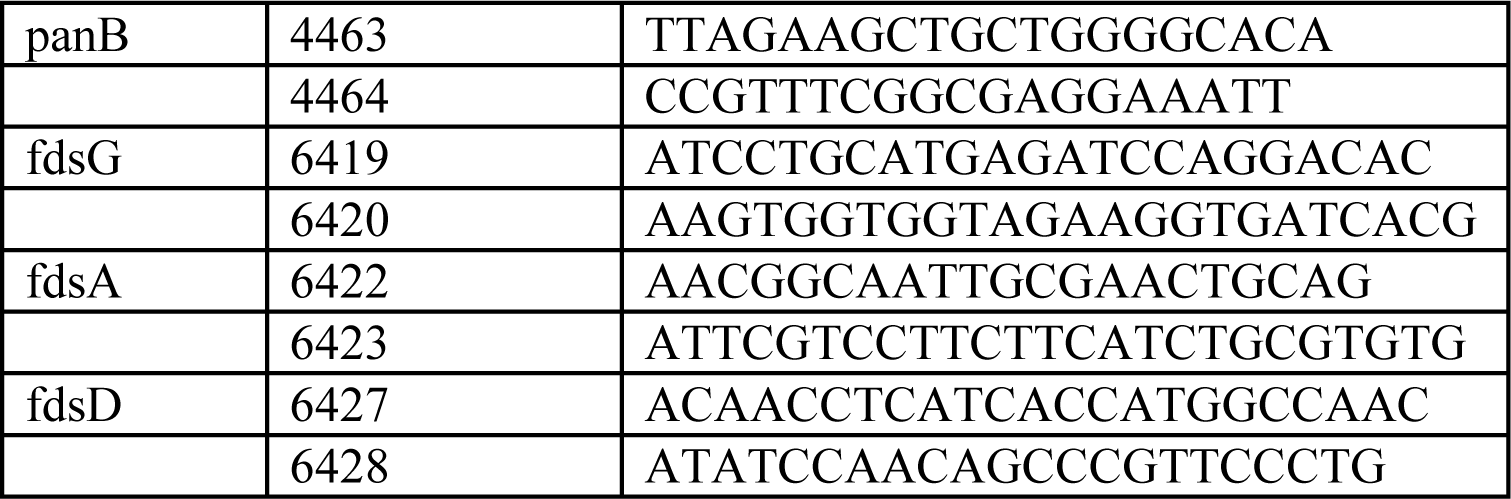

---

**Supplementary Table 1.** Mutations fixed in the genome of the evolved strains.

| UOF1 | chromosomal position* | mutation type | mutation event | mutated gene | intragenic position | amino acid change | intergenic mutation (distance to the flanking genes) | mutations identified in the indicated strain (•) |  |  |
| --- | --- | --- | --- | --- | --- | --- | --- | --- | --- | --- |
|  |  |  |  |  |  |  |  | **G5823** | **G5824** | **G5825** |
|  | 953670 | A/C | SNP | *focA* | 20 | F7C |  | • | • | • |
|  | 1029182 | C/A | SNP |  |  |  | yccW/yccX (-77/-105) | • | • | • |
|  | 1798380 | C/T | SNP | *infC* | 283 | E95K |  | • | • | • |
| UOF2 | chromosomal position* | mutation type | mutation event | mutated gene | intragenic position | amino acid change | intergenic mutation (distance to the flanking genes) | mutations identified in the indicated strain (•) |  |  |
|  |  |  |  |  |  |  |  | **G5848** | **G5849** | **G5850** |
|  | 88110 | A/G | SNP | *cra* | 83 | Y28C |  |  |  | • |
|  | 953401 | C/T | SNP | *focA* | 289 | V97I |  | • | • |  |
|  | 953670 | A/C | SNP | *focA* | 20 | F7C |  |  |  | • |
|  | 1785132 | G/A | SNP | *pps* | 5 | S2F |  | • | • |  |
|  | 2533860 | G/T | SNP | *crr* | 5 | G2V |  | • | • |  |
|  | 4159247 | T/G | SNP | *fabR* | 158 | L53W |  | • | • |  |

